# Pneumatically Controlled Non-Equibiaxial Cell Stretching Device with Live-cell Imaging

**DOI:** 10.1101/2023.06.15.545174

**Authors:** Jue Wang, Aritra Chatterjee, Clarisse Zigan, Maya Alborn, Deva D. Chan, Alex Chortos

## Abstract

**Objective:** Adherent cell behavior is influenced by a complex interplay of factors, including chemical and mechanical signals. *In vitro* experiments that mimic the mechanical environment experienced by cells *in vivo* are crucial for understanding cellular behavior and the progression of disease. In this study, we developed and validated a low-cost pneumatically-controlled cell stretcher with independent control of strain in two directions of a membrane, enabling unequal biaxial stretching and realtime microscopy during actuation.

**Methods:** The stretching was achieved by two independent pneumatic channels controlled by electrical signals. We used finite element simulations to compute the membrane’s strain field and particle tracking algorithms based on image processing techniques to validate the strain fields and measure the cell orientation and morphology.

**Results:** The device can supply uniaxial, equibiaxial, and unequal biaxial stretching up to 15% strain in each direction at a frequency of 1Hz, with a strain measurement error of less than 1%. Through live cell imaging, we determined that distinct stretching patterns elicited differing responses and alterations in cell orientation and morphology, particularly in terms of cell length and area.

**Conclusion:** The device successfully provides a large, uniform, and variable strain field for cell experiments, while also enabling real-time, live cell imaging.

**Significance:** This scalable, low-cost platform provides mechanical stimulation to cell cultures by independently controlling strains in two directions. This could contribute to a deeper understanding of cellular response to biorealistic strains and could be useful for future *in vitro* drug testing platforms.

## I. Introduction

A DHERENT Cell behavior is governed by an interplay of internal and external factors that include chemical and mechanical signals. These stimuli modulate cellular processes such as metabolism and extracellular matrix remodeling. In tissues such as lungs, arteries, cartilage and the musculoskeletal system, mechanical stimuli play an important role in regulating structure and function [1]. The biological response is highly dependent on the parameters of the mechanical stimulation (e.g., uniaxial vs biaxial, strain magnitude, frequency). The ability to mimic the complex mechanical stimuli is crucial to understanding mechanobiology in healthy tissues and diseases such as cardiac fibrosis [2]–[5], idiopathic pulmonary fibrosis [6], [7], as well as orthopedic [8] and connective tissue diseases [9].

Many cell stretching devices employ electrical actuators such as stepper motors [10]–[12], DC motors [13], voice coil motors [14], dielectric elastomer actuators (DEAs) [15] and shape memory alloys [16]. Commercial entities like Strex Inc. and CellScale also utilize motor-controlled actuation in their products. These electromechanical systems can have high precision and can include feedback. However, they are typically costly, limiting their use in large scale experiments. Furthermore, electrothermal heat generation can significantly affect temperature-sensitive cell culture experiments. Pneumatic actuators [17]–[20] have lower construction and maintenance costs and smoother motion due to continuous air flow. In addition, pneumatic systems pose a reduced risk of temperature fluctuations in sensitive incubator environments and a lower risk of contamination since the mechanical components do not come into direct contact with the cell culture apparatus. Some pneumatic systems [17], [21], including commercial products from FlexCell, require lubricants at the bottom of the membrane. These lubricants may interact with or seep into stretching membranes, affecting the behavior of cells. [22]

Regarding the functionality of cell stretching devices, uniaxial [11], [22], equibiaxial, and equiaxial [23]–[26] stretching methods are predominantly employed due to their relatively simple actuator designs. However, some tissues experience anisotropic mechanical cues. For example, some regions of the heart experience nearly uniaxial strains, some experience nearly equiaxial strains, while most exhibit non-equibiaxial strains. [27] Consequently, non-equibiaxial cell stretching devices provide a platform to facilitate a deeper understanding of cellular behavior relevant to these tissues and disease states. [28]

Currently, few studies focus on non-equibiaxial stretching. Wong et. al [28] and Hu et. al [29] proposed motor-driven non-equibiaxial stretching devices, but these systems preclude livecell imaging. Tremblay et al. [30] introduced a pneumatic cell stretching device with two independent channels but did not demonstrate non-equibiaxial stretching; the small deformation chamber also hindered the generation of a uniform strain area. Essential design features for a versatile stretching device that can be applied to various cell study systems include independent control of amplitude and frequency in two orthogonal directions, maintenance of strain homogeneity in thin planar materials, and compatibility with high-resolution optical imaging for real-time data acquisition. To address the limitations of currently available designs, in this work, we have developed and validated a pneumatically controlled cell stretcher with independent control of strain in two directions of a microscopy-compatible membrane. In-plane actuation allows large area uniform non-equibiaxial stretching and real-time microscopy during actuation. Our device is sized similarly to 60-mm petri dishes, making it compact and lightweight. Furthermore, the pneumatic control system can accommodate control of multiple parallel devices under identical actuation conditions.

## II. Materials & Methods

### A. Design and fabrication of bi-directional cell stretcher

The design of the bi-directional cell stretcher is depicted in Fig. 1(a). It consists of 4 main parts: stretcher base, vacuum cavity, stretching membrane, and stickers. The design of the base enables mounting onto an inverted microscope stage for live-cell imaging and connections with the pneumatic system. Four quarter-circular vacuum cavities generate a bi-directional stretch, with two opposing cavities connected to a single pneumatic channel to ensure uniform strain in each direction. Vacuum pressure generates shrinkage of cavities’ inner shell, stretching the membrane attached to the shell’s bottom. Rigid stickers at the edges of the membrane guide the deformation of the membrane to enable uniform strains. The working area is defined by the size of stickers, which is a 22*mm* × 22*mm* square as shown in Fig. 1(b). We used spray paint to create a speckle pattern to track strain under the microscope. Fig. 1(c) & (d) depicts the deformation on the membrane when we applied non-equibiaxial stretching, which demonstrates the basic functionality of the stretcher in this paper.

**Fig. 1.**
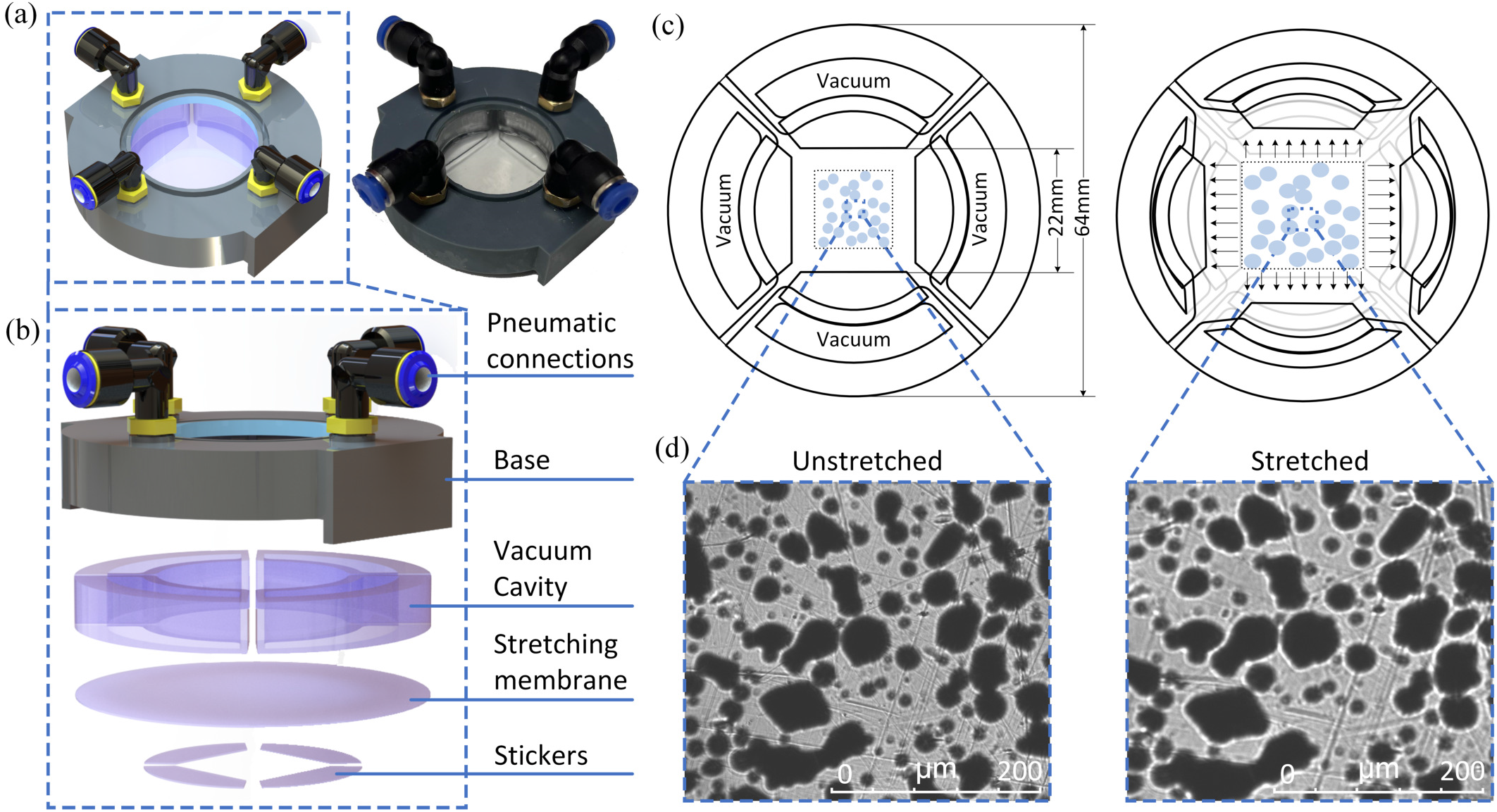
The design and principle of bi-directional cell stretcher. (a) The 3D model and actual device of bi-directional cell stretcher. (b) The explosion diagram of bi-directional cell stretcher. (c) The dimension and principle of bi-directional cell stretcher. (d) Stretching of a speckle pattern.

The base is made by stereolithography (SLA) 3D printing using a Phrozen 8k printer with 8k Aqua Gray resin and cured in an oven at 70^°^*C* for 24 hours to ensure biocompatibility. [31] Both cavities and membrane are made of polydimethylsiloxane (PDMS) Sylgard 184 (Dow Corning, USA) with base to cross-linker ratio 17.5:1 and fabricated using molds which are printed with the same printer. The mold has two parts: the middle part defines the vacuum cavity while the outer part of the mold defines the outer shell of the cavities. The Sylgard 184 was mixed using a centrifugal mixer for 4 min and was subsequently centrifuged for 30*s* to remove the bubbles. After pouring the Sylgard 184 into the mold, the Sylgard 184 was degassed for 30*min* in a vacuum chamber. Following this, the molds with uncured PDMS 184 are put into the oven at 60^°^*C* for 24 hours. The membranes are fabricated by blade coating uncured Sylgard 184 to a uniform thickness of 0.4*mm* onto glass coated with non-stick Bytac coating (Cole-Parmer Bytac D1069324 Surface Protector, Fisher). Following this, the membrane was cured in an oven at 60^°^*C* for 24 hours. The rigid stickers laser cut from a 1*/*32^′′^-thick acrylic sheet. The inner and outer diameter of the quarter-circular cavities are 34*mm* and 64*mm*, respectively. The thickness of the stretching membrane attached to the cavities is 0.4*mm*.

To ensure the whole device is biocompatible, uncured Sylgard 184 was used to adhere the individual components (base, vacuum cavities, stretching membrane and stickers) together. The entire device was placed in an oven at 60*C* for 24 hours to cure the Sylgard 184 adhesive. (Fig. 2)

**Fig. 2.**
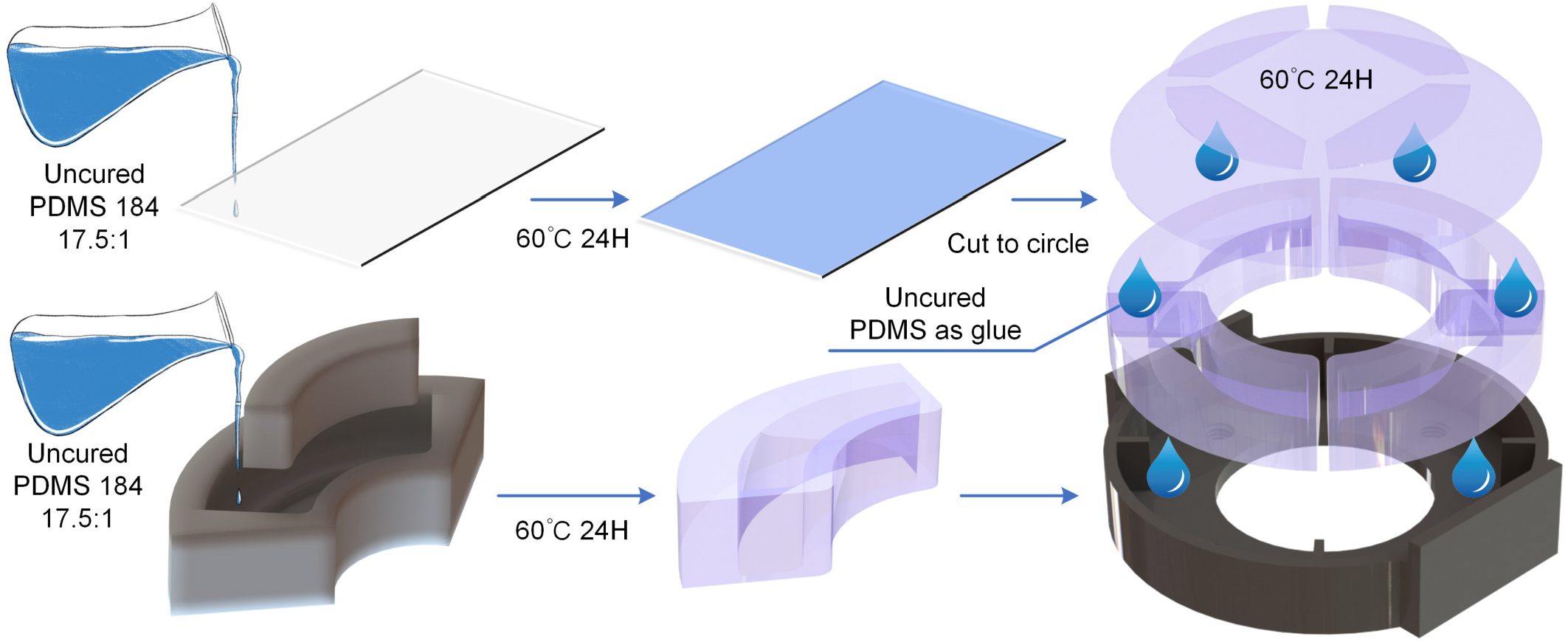
The fabrication process of bi-directional cell-stretcher.

All materials in contact with culture media were tested separately for biocompatibility. We also tested for biocompatibility of the entire system by performing live-dead staining on 3T3 fibroblasts cultured using DMEM +10% FBS+1% Pen Strep inside the cell stretcher system for 24 hours under standard culture conditions (37*C*, 95% humidity, 5% *CO*_2_)

### B. Dual-channel pneumatic control system

Negative pressure was modulated using a vacuum generator (Festo VAD 1/4) that was driven by positive pressure from 0 to 100 psi and controlled using a Alicat Electro-pneumatic Transducer (PCD-series). 0 − 5*V* analog control signals for the Alicat were generated using digital-to-analog converters (MPC4725) controlled using an Arduino (Mega 2560). The communication between the Arduino and MPC4725 used an I2C interface. However, controlling the two actuation directions independently requires two channel inputs while the Arduino has only one hardware I2C channel. We used software I2C to simulate two channels to communicate with two MPC4725 converters. (Fig. 3)

**Fig. 3.**
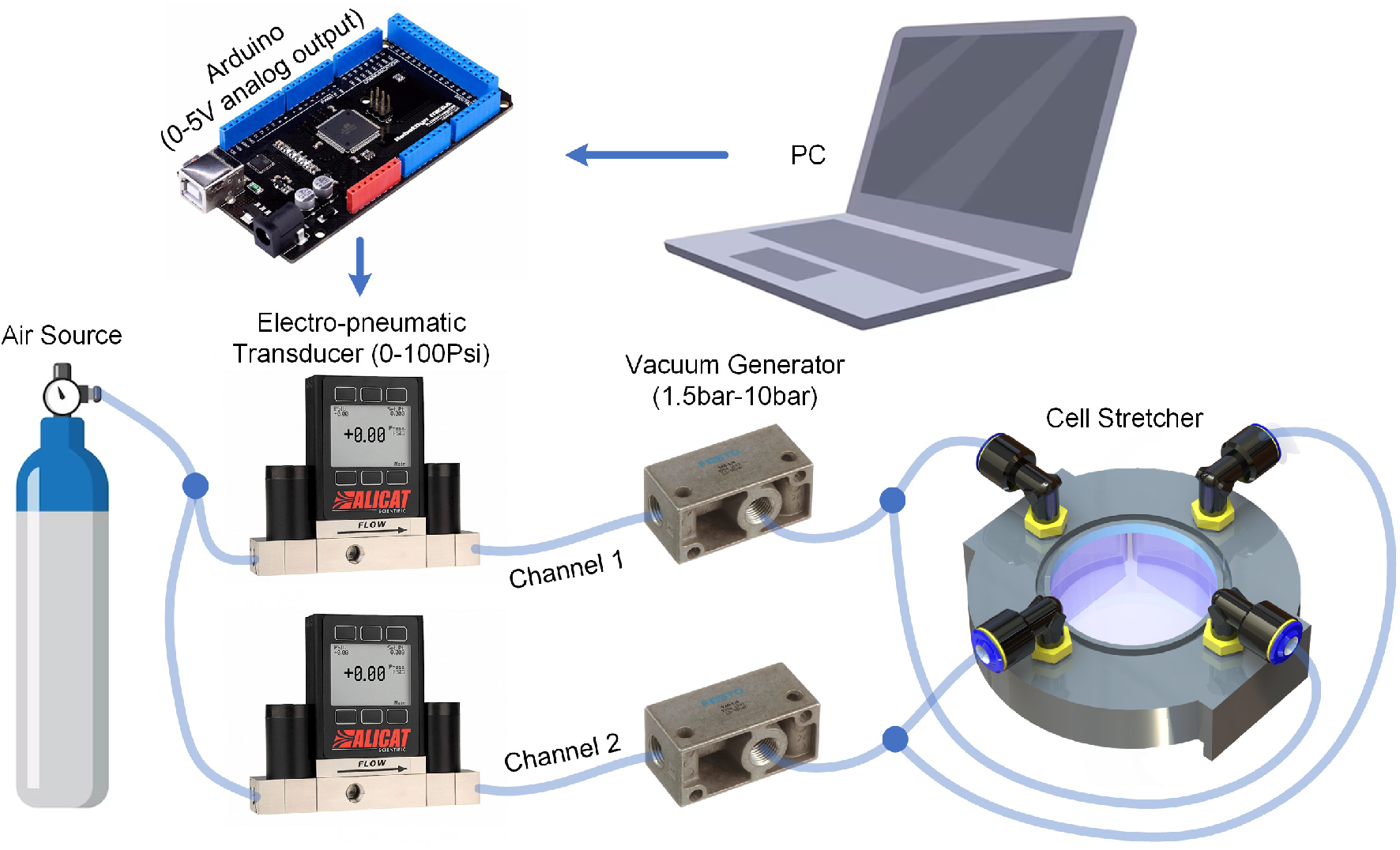
The control system of bi-directional cell-stretcher.

### C. Finite element (FE) simulation of cell stretcher

Initially, we examined the material properties by conducting a tensile test. Dogbone samples with a 17.5:1 ratio of Sylgard 184 were mounted onto an Electroforce 5500 system. The samples were loaded at 10 mm/min until failure. In total, five samples were tested. The Young’s modulus extracted from the linear strain region (10%) was found to be 303.26±11.56*kPa*. In addition, the elongation at break for the samples was 240%.

To predict the strain fields on the membrane under different vacuum inputs, finite element (FE) simulations were performed using COMSOL (COMSOL, Inc., USA). The 3D model was made by Solidworks and imported into COMSOL for analysis using the Solid Mechanics module. PDMS was modeled as a linear elastic material with Young’s Modulus of 303*kPa* as derived above and Poisson’s ratio of 0.43. The base made of SLA resin was assigned a Young’s Modulus of 2.2*GPa* and Poisson’s ratio of 0.4. The acrylic stickers were assigned a Young’s Modulus of 2.5*GPa* and Poisson’s ratio of 0.4. In the simulation, the vacuum pressure is converted to the negative pressure applied to the inner shell of cavities. The mesh is defined as ‘Normal’ size.

### D. Quantification of strain fields in stretched substrates

Strain quantification experiments were performed to determine the region of uniform strain in the stretched substrates and calibrate the pneumatic loading system. Specifically, fiduciary markers on the PDMS membranes were tracked using a video camera, under different stretch amplitudes and frequencies in both uniaxial and biaxial directions. From the acquired images, marker displacements were quantified using MATLAB (R2021a) between the reference and deformed configurations using particle tracking. A strain interpolation algorithm was implemented to quantify the in-plane Green-Lagrange strains from the marker displacements. gradients [32].

### E. Substrate ligand coating and cell culture

The PDMS membrane based stretching device was sterilized and the substrates were cleaned by ultrasonication using 70% ethanol followed by distilled water for 20 minutes each. The entire surface of the PDMS substrates were then coated with 3 ml of 80 μg/mL collagen solution (Type I Collagen Solution, 4mg/ml (Rat Tail)) at 37^°^*C* for 1 hour to facilitate cell attachment. Following which, the excess solution was then aspirated and the substrates were washed with PBS prior to seeding the cells. GFP expressing NIH-3T3 fibroblast cells were cultured on the coated PDMS substrates using Dulbecco’s Modified Eagle Medium (DMEM) supplemented with 10% (v/v) fetal bovine serum (FBS) and 1% (v/v) penicillin/streptomycin in a sterile incubator with a humidified atmosphere containing 5% *CO*_2_ at 37^°^*C*. A cell seeding density of 5*x*10^4^*cells/mL* was used for the stretching experiments. Cells were incubated overnight on the stretching device to allow attachment to the substrate prior to the stretching experiments.

### F. Live-cell imaging under stretch

The pneumatically controlled stretching device with cells cultured on the PDMS membrane was mounted on a Leica DMI6000b epifluorescence microscope for real-time live imaging of cells under stretch. The microscope was housed inside a caged incubator (Pecon) equipped with temperature and *CO*_2_ controllers, that helped maintain humidified atmosphere at 5% *CO*_2_ and 37^°^*C* required for prolonged cell viability. Cells were imaged with a 20x objective using brightfield and fluorescence microscopy to quantify changes in cell morphology under stretch. To augment the features and reduce background interference in the phase contrast images, fundamental post-processing steps were implemented in ImageJ (NIH). These included Fast Fourier Transformation (FFT) bandpass filtering, image sharpening, and contrast enhancement. The derived binary image was then employed for cell orientation analysis. Cell stretching experiments were performed at 1*Hz* for 3 hours for all sinusoidal strain combinations. Live-cell images were captured at 15-minute intervals.The loading briefly paused for a few seconds during capturing images, ensuring that each image taken was in a state without loading. At the beginning and end of each stretching process, we evaluated cell orientation in 5 different regions of interest (ROIs).

## III. Results

### A. System characterization of bi-directional cell stretcher

The in-plane membrane strains were quantified by tracking fiducial markers on the PDMS using a particle tracking algorithm. To characterize the strain-pressure relationship, we first applied a continuously varying pressure in the x and y directions separately as shown in Fig. 4(a). The result shows the change in strain is relatively linear, especially in the latter half when pressure decreases linearly. The strain in both directions reach a maximum of 15% strain at 48*psi*. The nearly identical pressure-strain relationship in the x and y directions reflects the uniformity of the stretcher. As shown in Fig. 4(b), we validated the stretcher actuation under sinusoidal loading (1*Hz*). The strain of the stretcher closely follows the sinusoidal pressure input. Therefore, in the following experiment, we chose sinusoidal loading as our input for cell experiments.

**Fig. 4.**
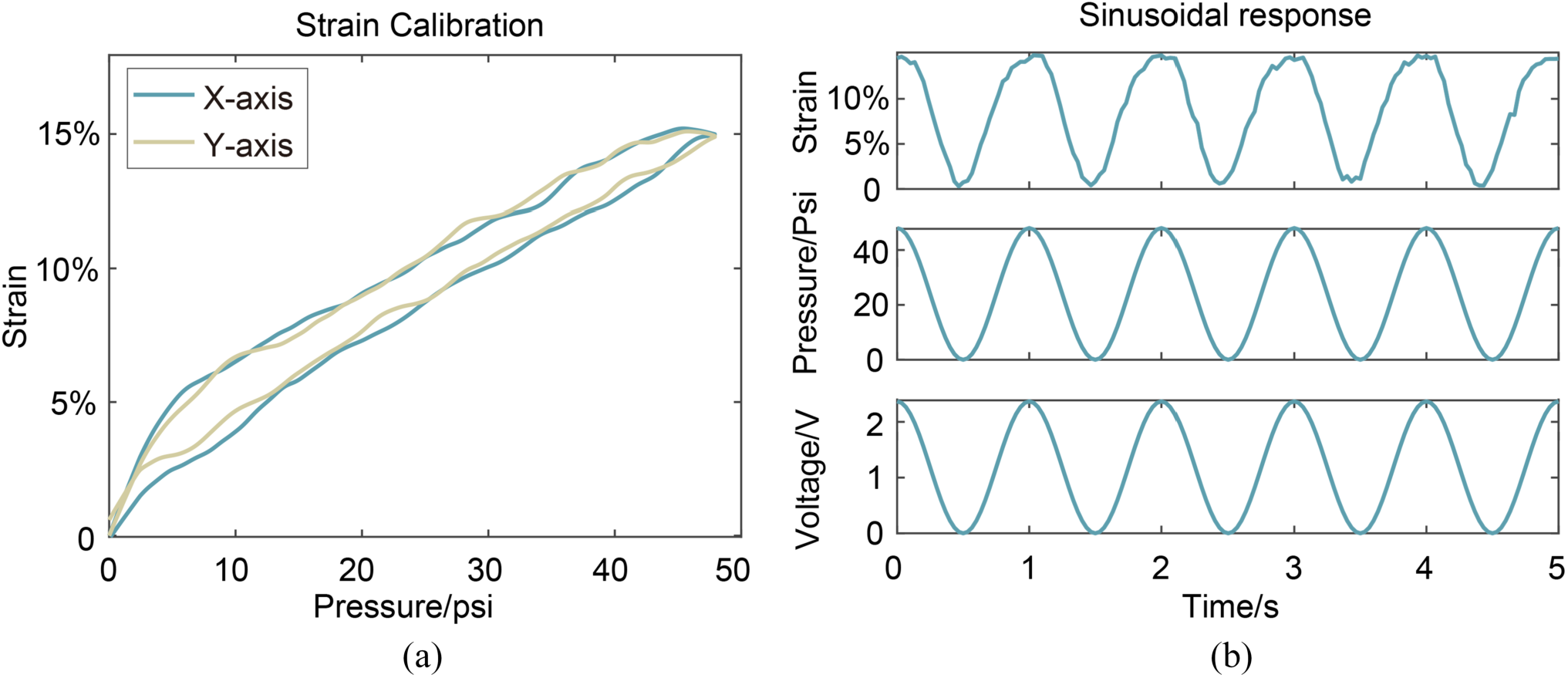
System characterization of cell stretcher. (a) X-axis and Y-axis strain under continuous varying pressure. (b) uniaxial strain under sinusoidal loading.

Next, we validated the functionalities of the stretcher under sinusoidal loading. We compared three cases: uniaxial, equibiaxial and non-equibiaxial loading by experimentally quantifying the in-plane strains in each case. In the uniaxial case, we provide one channel of sinusoidal loading to only stretch the x-axis. The maximum strain amplitudes are calibrated to 5%, 10% and 15% with 16*psi*, 32*psi* and 48*psi*, respectively. From Fig. 5(a), we observe that the device has a stable and repeatable sinusoidal response to the loading input, but the uniaxial stretching in x-axis can cause a heterophase strain change (20% of the x-axis strain) in y-axis. This Poisson effect has been reported in other uniaxial stretcher work [30], [32]. However, for our bi-directional stretcher, the actuation in the other direction is able to compensate for the Poisson effect. FE simulations shown in Fig. 5(b) show more than 70% of the central area of the membrane exhibits uniform strain with average E11 (x-axis strain) of 5.1%, 10.2%, 15.3%, respectively and average E22 (y-axis strain) −1.2%, −2.5%, −3.8%, respectively, which corresponds well with the experimental measurements. In the equibiaxial case, we provide two channels of homophase sinusoidal loading. As shown in Fig. 5(c), when applying 30*psi* − 29*psi* in both channels, the stretcher can obtain exactly 5% strain in both directions. Similarly, when applying 49*psi* − 48*psi* and 65*psi* − 64*psi*, equibiaxial strains of 7.5% and 10% could be obtained. The slight difference in the calibrated pressures may be caused by small variations in the pneumatic controls for the two independent channels. Example sources of these errors could include variations in the length of the tube used to connect the stretcher to the vacuum source or the calibration error of the pressure controller. The unique capability of this stretcher to generate strains in two directions independently is demonstrated using 3 example cases of strains in the x-y directions: 2.5% − 7.5%, 5% − 10%, and 10% − 5%. For each example, the corresponding input pressure is 24*psi* − 48*psi*, 37*psi* − 53*psi*, and 55*psi* − 35*psi*, respectively. Fig. 5(d) & (f) depicts the FE simulation results that are under the same pressure conditions of bi-directional stretching. The results agree with their relevant experimental measurements within 5% error in both directions.

**Fig. 5.**
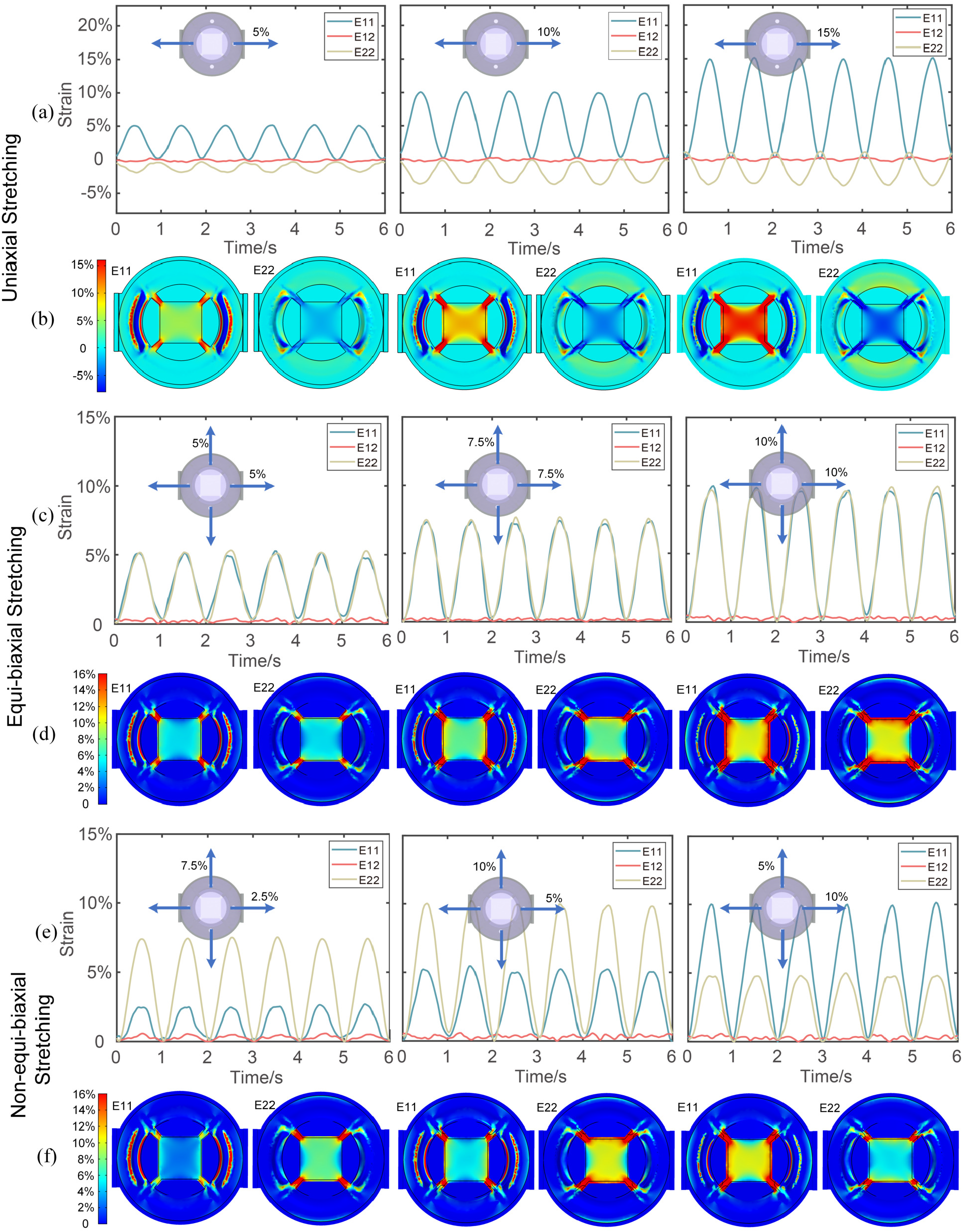
The demonstration of the functionalities of bi-directional cell stretcher. (a)-(b) Experimental results and simulation results of uniaxial stretching of 5%, 10%, and 15% under 1*Hz* sinusoidal loading. The relevant pressure input is 16*psi*, 32*psi*, and 48*psi*. (c)-(d) Experimental results and simulation results of equibiaxial stretching of 5%, 7.5%, and 10% under 1*Hz* sinusoidal loading. The relevant pressure input is 30*psi* − 29*psi*, 49*psi* − 48*psi* and 65*psi* − 64*psi*. (e)-(f) Experimental results and simulation results of non-equibiaxial stretching of 2.5% − 7.5%, 5% − 10%, and 10% − 5% under 1*Hz* sinusoidal loading. The relevant pressure input is 24*psi* − 48*psi*, 37*psi* − 53*psi*, and 55*psi* − 35*psi*.

### B. Live-cell images on static-stretched membrane

A significant advantage of the proposed cell stretching device is the ability to capture high-quality live-cell images through the stretchable PDMS membrane. To capture stable images, we stretched the membrane uniaxially and bi-directionally to 10% static strain and recorded live-cell images every 20 min (Fig. 6). All images were captured in real-time directly from the microscope without any postprocessing. Compared to the control (no strain), the 0 min (10% strain) images indicate that the adherent 3T3 cells are sensitive to the applied strain on the underlying membrane which affects their relative position and morphology. This stretching process generates internal stress within the cells, which progressively modifies their shape. In the uniaxial case, the stress experienced by the cells is oriented in one direction, causing cells aligned with the stretching direction to initially become thinner. Over time, this internal stress prompts partial detachment of certain cells from the membrane, culminating in a rounded morphology. In the bi-directional case, since the internal stress is uniformly distributed in both directions, cells do not experience extensive detachment as in the former case. Instead, only a few individual cells exhibit a reduced adhesion area with the membrane.

**Fig. 6.**
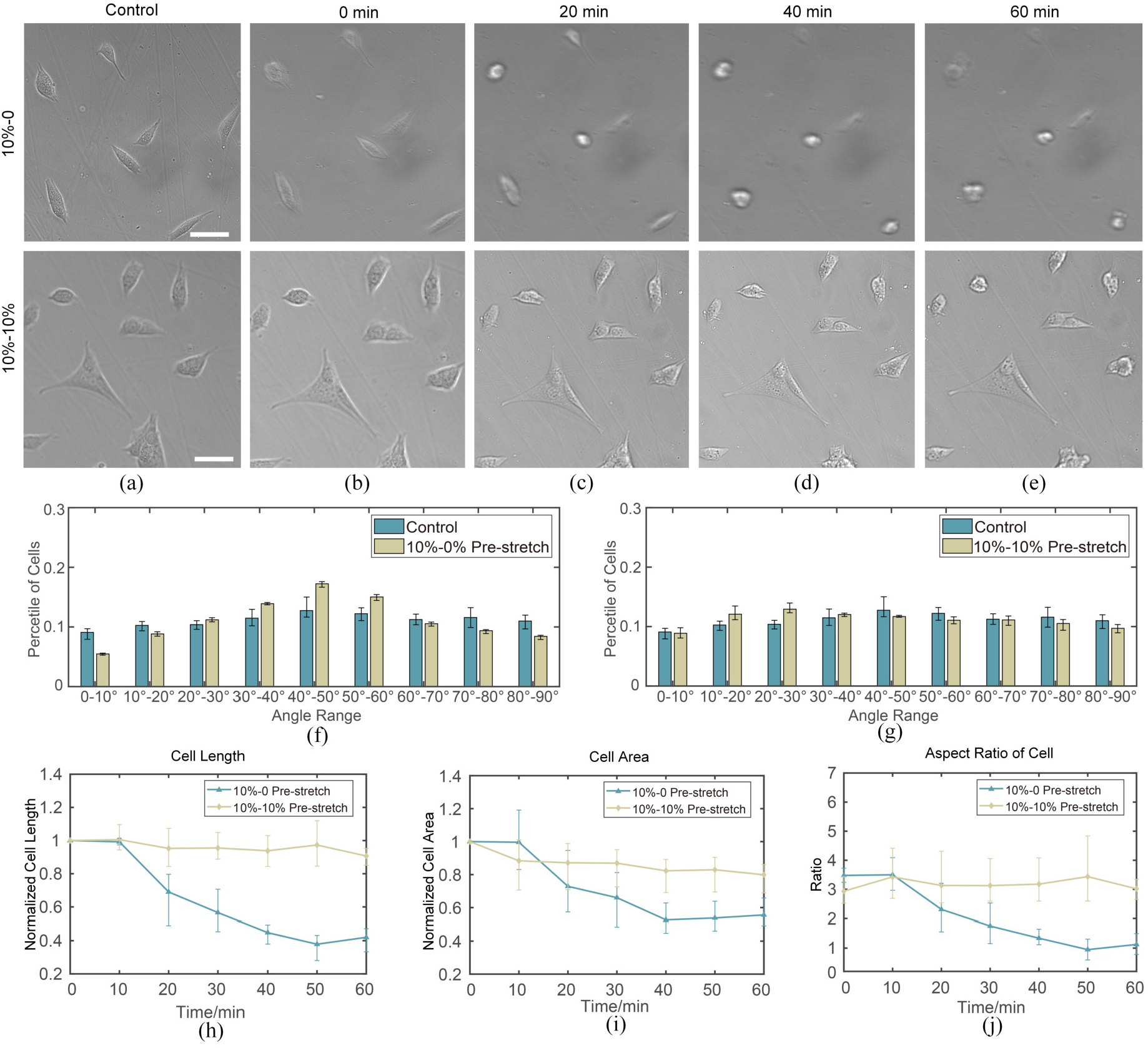
Cells subjected to static stretch under uniaxial and bi-directional loading shown with bright field microscopy images (unprocessed cell images). (a)-(e) Live-cell images of unstretched membrane (Control) and stretched membrane every 20min. (f) Cell orientation changes after applying 1 hour static stretch. (g)-(i) depicts the changes in single cell morphology, where (g) shows the normalized length change of cells and (h) shows the area change of cells under two different stretching conditions. (i) displays the aspect ratio (ratio of the major to minor axes for individual cells, with the assumption that cells exhibit an elliptical shape). The white scale bar in (a)-(e) refers to 50*μm*.

We performed a quantitative assessment of cellular orientation and morphology change, as illustrated in Fig. 6(f). The application of 1 hour static uniaxial stretching resulted in not only partial cell detachment but also a gradual realignment of the initially uniformly distributed cells in the orthogonal direction, leading to a major angle of approximately 45 degrees. In contrast, the 1 hour static bi-directional stretching demonstrated minimal influence on cell orientation. We employed single-cell tracking to quantify changes in cellular morphology. Initially, we monitored alterations in the length and area of individual cells. (Fig. 6(g) & (h)) Subsequently, by assuming an elliptical cell shape, we calculated the ratio between the major and minor axes for more intuitive description. (Fig. 6(i)) uniaxial stretching leads to a reduction in cell length and causes detached cells to adopt a more circular shape. In contrast, bi-directional static stretching does not induce any significant morphological changes in cells. Notably, while a subset of cells becomes thinner, another subset appears to become more rounded.

### C. Cellular orientation and morphology change under cyclic stretching with different strain conditions

To verify the functionality of the device in cell experiments, we quantified cell alignment under four distinct stretching conditions: 10% − 0% uniaxial stretching, 10% − 10% equibiaxial stretching, 10%−5% and 5%−10% non-equibiaxial stretching.

As demonstrated in Fig. 7(a), uniaxial stretching results in cellular realignment perpendicular to the stretch direction. While equibiaxial stretching can induce similar cellular realignment with less pronounced realignment magnitude (Fig. 7(b)). non-equibiaxial cell stretching is also capable of inducing notable alterations in cell orientation. For the 10% − 5% scenario, the predominant cell angles span from 50 to 70 degrees, attributable to the higher strain experienced in the x-direction. In contrast, the 5%−10% situation exhibits primarilay cellular orientation angles between 10 and 30 degrees, resulting from the higher strain present in the y-direction.

**Fig. 7.**
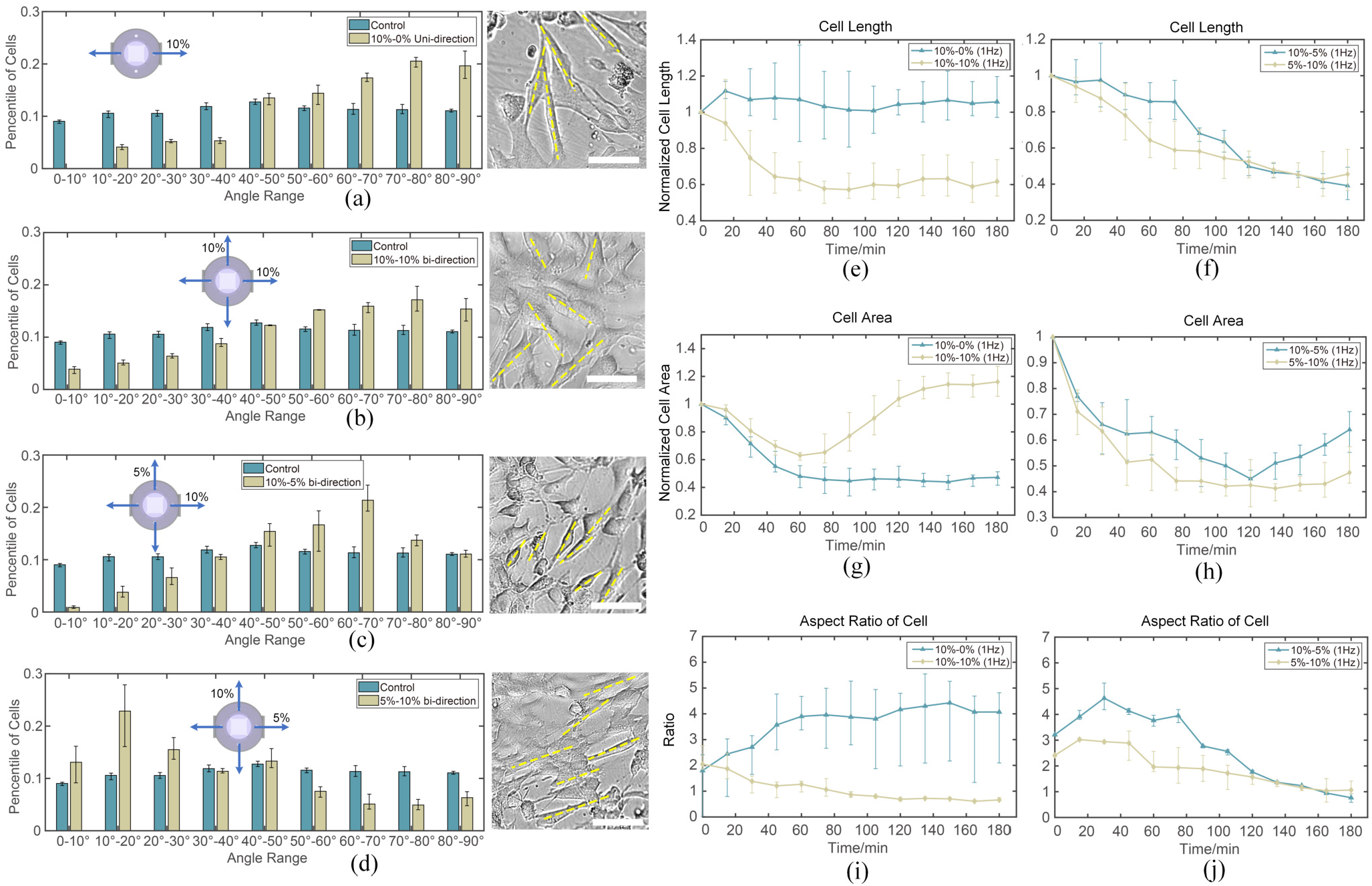
Changes in cell orientation and morphology after 3 hours of cyclic stretching. (a) Cell orientation change after 3 hours 10% − 0 uniaxial stretching. (b) Cell orientation change after 3 hours 10% − 10% equibiaxial stretching. (c) Cell orientation change after 3 hours 10% − 5% nonequibiaxial stretching. (d) Cell orientation change after 3 hours 5% − 10% non-equibiaxial stretching. The angle under consideration represents the inclination relative to the x-axis (horizontal). 0^°^ denotes alignment parallel to the x-axis, whereas 90^°^ indicates orientation perpendicular to the x-axis. (e)-(j) depicts the morphology change of single cells, where (e)-(f) shows the normalized length change of cells and (g)-(h) shows the normalized area change of cells under four stretching conditions. (i)-(j) displays the aspect ratio (ratio of the major to minor axes) for individual cells, with the assumption that cells exhibit an elliptical shape. In (a)-(d), we show the bright field microscopy images (unprocessed cell images) after 3 hours stretching and have used yellow dashed lines to indicate the orientation of the cells. The white scale bar in (a)-(d) refers to 40*μm*.

Additionally, utilizing the live-cell images acquired during the experiment, we monitored the area and length alterations of 5 individual cells in each case. Here, we first transform the image series collected during the cell stretching into a video and then used the Direct Linear Transformation (DLT) digitizing tool implemented in Matlab. It can automatically track and analyze objects in video sequences (Developed by Ty Hedrick from UNC). [33] When selecting tracking cells, we chose five cells with initial orientations closely aligned with the y-axis. Within the DLTdv8a, the two vertices of the cells’ longer edges were selected for tracking. If the auto-track is interrupted due to sudden shape change of cells between frames or insufficient image clarity, manual selection of the tracking points is employed. From this data, we obtained the variation in cell length over time. (Fig. 7) Subsequently, we monitor the alterations in the area of these cells by initially delineating the cellular contours and subsequently quantifying the pixel counts associated with respective shapes. (Fig. 7) To more effectively illustrate the morphology change of the cells from the data, we posited that the cell’s overall shape is elliptical.

Then, we derived the ratio of the major and minor axes by the data of the cell’s length and area. Fig. 7(e) & (f) demonstrates that uniaxial stretching leads to an increased elongation of the cell and a decrease in overall cell area (Fig. 7(g)), while bi-directional stretching results in a more rounded cellular morphology (Fig. 7(g)). We also observe that the changes in cell length and area under the application of non-equibiaxial stretch follow similar trends irrespective of the direction of applied stretches (Fig. 7(f) & (h)). Finally, the changes in cellular aspect ratio corroborate with the changes in cell length and area with cells becoming more elliptical under uniaxial stretch as compared to equibiaxial stretch (Fig. 7(i)). The aspect ratio changes under anisotropic stretch show analogous responses for both 10%5% and 5% − 10% stretch.

## IV. Discussion

In this study, we describe a custom mechanobiological platform that enables measuring cellular responses to non-equibiaxial strains over a range of frequencies and strain magnitudes. This first manifestation of the platform described in this manuscript has been designed with a large cell growth area (2*cm* × 2*cm*) for cell attachment and visualization. However, our design can be scaled to smaller or larger dimensions based on the needs of the research. Compared with microscale cell growth area [30], [34], larger culture areas provide a more homogenous strain field that could provide more reliable and robust results. Furthermore, our scalable cell culture area could be advantageous for studies necessitating a high cell count or the collection of multiple samples from a single culture.

While this work was demonstrated with 2D cell cultures, it is widely accepted that 3D cell cultures more closely replicate the *in vivo* environment. [35], [36] To theoretically evaluate the ability to facilitate 3D cell culture, we have used finite element simulations to determine the impact of an appropriate thickness of collagen (30*μm*) on membrane stretching. The presence of a cell-laden collagen gel with a Young’s modulus of 15*kPa* [37] does not significantly affect the membrane’s stretching performance because its stiffness is significantly lower than that of the membrane. The results revealed that, under the same conditions, a membrane with 30 μm collagen stretched to a 10% strain while a membrane without collagen reached a 10.4% strain. Consequently, the presence of collagen only introduces discrepancy of around 4%, which can be corrected in the calibration process.

The last several decades of mechanobiology research have emphasized the breadth of parameters that affect cell growth, including the modulus, porosity, and viscoelasticity of the extracellular environment in addition to chemical stimuli. Increasing evidence shows that these factors can be interdependent, necessitating the performance of combinatorial measurements to elucidate their effects. [38], [39] These combinatorial measurements require parallel experiments on many conditions simultaneously, which could be enabled by the low cost of our system.

The material cost of each stretcher priced at approximately $10 (including the 3D print resin and PDMS). Following the completion of the 3D-printed mold, the device can be affordably mass-produced. As a single control system can simultaneously manage more than 10 stretchers, its control system cost is relatively low compared to motor-driven cell stretchers, each of which requires an individual motor for operation. Concurrently, upon the conclusion of cell culturing, reusability can be attained through membrane replacement. Provided that the membrane’s thickness remains consistent and uniform, the error resulting from membrane substitution can be maintained below 3%. Moreover, as all chambers are generated from an identical mold, the variability between stretchers is less than 5%, primarily attributable to manual assembly in laboratory settings. It is anticipated that the error will decrease further with the implementation of industrial-scale mass production.

The main cost of the system is the pneumatic controls. Our system exhibits an average pumping speed of 30*L/min*, while the individual chamber volume is approximately 1*cm*^3^. Consequently, the time delay between operating 10 stretchers concurrently versus a single stretcher is less than 0.04 seconds, which is much smaller than the rate at which we actuate the devices (1*Hz*). This indicates that a single pneumatic control system can efficiently actuate over 10 stretchers simultaneously, which allows for higher-throughput studies.

## V. Conclusion

In this work, we present a novel cell stretching system that enables independent control of orthogonal in-plane strains on a deformable membrane, through which live-cell imaging is possible. Uniaxial and biaxial cell stretching devices are useful to quantify cellular response to complex mechanical stimulation, which could give insights into tissue development or disease progression. Our strain calibration experiments show that our custom stretching device can provide controllable multiaxial mechanical stimuli. Due to the in-plane deformation of the cell culture membrane, real-time imaging was used to track the cell size and orientation as a function of static and cyclic strains. Significant differences were observed in the cell aspect ratio and alignment when using the non-equibiaxial strains enabled by our device compared to the uniaxial and biaxial strains that are commonly used. We believe our device will be useful for conducting a range of mechanobiological studies, providing useful insights into cell behavior in the context of disease diagnosis and development.

## Notes

This work was supported in part by the National Science Foundation under Grant 2149946 to Deva D. Chan and in part by the Purdue startup funding to Deva D. Chan and Alex Chortos.

### Competing Interest Statement

The authors have declared no competing interest.

